# Shared Characteristics of Intrinsic Connectivity Networks Underlying Interoceptive Awareness and Empathy

**DOI:** 10.1101/2020.04.30.070490

**Authors:** T. Stoica, B.E. Depue

## Abstract

Awareness of internal bodily sensations (interoception, IA) and its connection to complex socioemotional phenomena like empathy have been postulated, yet its neural basis remains poorly understood. The present fMRI study employs independent component analysis (ICA) to investigate whether the Cognitive or Affective facets of empathy and IA share resting state network connectivity and/or variability (SD_BOLD_). Healthy participants viewed an abstract movie demonstrated to evoke strong connectivity in resting state brain networks (*InScapes*), and resultant connectivity and variability data was correlated with self-reported empathy and IA questionnaires. We demonstrate a bidirectional behavioral and neurobiological relationship between empathy and IA, depending on the type of empathy interrogated: Affective empathy and IA share both connectivity and variability, while Cognitive empathy and IA only share variability. Specifically, increased connectivity in the right inferior frontal operculum (rIFO) of a larger attention network was associated with increased vicarious experience but decreased awareness of inner body sensations. Furthermore, increased variability between brain regions of an interoceptive network was related to increased sensitivity to internal sensations along with discomfort alleviation arising from witnessing another’s distress. Finally, increased variability between brain regions subserving a mentalizing network related to not only an improved ability to take someone’s perspective, but also a better sense of mind-body interconnectedness. Overall, these findings suggest that the awareness of one’s own internal body changes (IA) is related to the ability to feel and understand another’s emotional state (empathy) and critically, that this relationship is not task-dependent, but is reflected in the brain’s resting state neuroarchitecture. Methodologically, this work highlights the importance of utilizing network variability as a complementary window alongside functional connectivity to better understand neurological phenomena. Our results may be beneficial in aiding diagnosis in clinical populations such as autism spectrum disorder, where participants may be unable to complete tasks or questionnaires due to the severity of their socioemotional symptoms.

## 1 Introduction

Internal body signals relative to emotion processing has been a topic of long-standing interest (Gurney 1884; Strack, Martin, and Stepper 1988), with more recent evidence highlighting an intriguing bidirectional relationship between sensations that arise internally and emotional phenomena (Lane 2008; Craig 2009; Cameron 2001; Damasio 2005). A proposed biological basis that may clarify this interplay is interoception, namely – the afferent processing of internal bodily signals that arise from visceral organs (Craig 2009; Cameron 2001; Wiens 2005; Johnson 2001; Cacioppo et al. 2000). Indeed, neuroimaging findings corroborate a substantial overlap between the neural substrates of emotion and interoception (Critchley and Garfinkel 2017; Adolfi et al. 2017), supporting the idea that the central monitoring and representation of internal bodily signals plays a fundamental role in emotion. However, the relationship between the emotions of *another* individual and one’s own body remains relatively unexplored.

Consider the ubiquitous scenario of listening to a close friend share a difficult moment. In order to accurately grasp the story from their perspective, you must map their internal sensations onto your emotional landscape, thereby creating a bridge towards proper understanding. This process hinges on two crucial socioemotional constructs: interoceptive awareness (IA) and empathy. IA represents the ability to accurately and consciously perceive changes in your internal bodily signals (Khalsa et al. 2018), while empathy refers to both vicariously experiencing and understanding the mental state of your friend (Davis 1980). Thus, empathy can be further fractioned into two interrelated facets: Affective and Cognitive empathy. Affective empathy is conceptualized as the automatic process of vicariously experiencing the emotional state of another (Baron-Cohen and Wheelwright 2004; Davis 1980), while Cognitive empathy describes the individual’s ability to accurately imagine another person’s perspective (Davis, 1980; Decety & Jackson, 2004; Lawrence et al., 2006). Hence, in the context of the anecdote above, genuine understanding arises when the listener is aware of their own internal bodily sensations while taking into account the speaker’s feelings and mental state. However, despite their importance in harmonious socioemotional exchange, it is currently unknown which specific facets of empathy dovetail with IA.

Neuroimaging research demonstrates that the neural substrates of empathy consistently overlap with the cortical regions involved in self-experience (Iacoboni 2005; Jackson, Meltzoff, and Decety 2005; Keysers et al. 2004; Wicker et al. 2003). For instance, during the experience of various aversive taste stimuli, activation of the anterior insula (AI) and inferior frontal operculum (IFO) were observed in both the observer and experiencer (Jabbi, Swart, and Keysers 2007). Similarly, observing others’ pain has been found to robustly activate the AI and anterior cingulate cortex (ACC) (Lieberman and Eisenberger 2009; Jackson, Meltzoff, and Decety 2005; Singer et al. 2004), regions associated with one’s own pain. One popular interpretation of such ‘shared representation’ between self and other’s experience (Decety and Sommerville 2003) posits that the brain represents others’ experiences in terms of the experiences of the self. To wit, one’s own internal state serves as a blueprint for understanding the experiences of others (Iacoboni 2009; Singer and Lamm 2009; Rizzolatti, Fogassi, and Gallese 2006). Although this view has been broadly (and often implicitly) accepted, the neurobiological details of this putative self-referential process for understanding others remain largely unclear.

This lacuna may exist due to the interacting, but only partially overlapping neural bases of empathy’s two facets (Fan et al. 2011). Affective empathy primarily elicits activations from regions implicated in rapid and prioritized processing of emotion signals, including: the amygdala, hypothalamus, orbitofrontal cortex (OFC) and AI (Decety et al. 2013). By comparison, Cognitive empathy, which shares similar neural networks with perspective-taking and mentalizing (Pardini and Nichelli 2009) additionally involves the superior temporal sulcus (STS), temporoparietal junction (TPJ), fusiform gyrus (FG), and medial prefrontal cortex (mPFC; (Saxe and Powell 2006)). To date, only one recent meta-analysis investigated convergent areas of activation between interoception, emotion and social processing (Adolfi et al. 2017). The results for the three domains converged in the AI, amygdala, right inferior frontal gyrus (rIFG), basal ganglia and medial anterior temporal lobe (mATL), ascribing particular importance to the fronto-temporo-insular nodes (Adolfi et al. 2017). The authors conclude co-activation of these regions may result in an evaluative association of the internal milieu, and in combination with external cues, leads to complex social behavior (i.e. empathy) (Adolfi et al. 2017).

Although this activation-based analysis revealed a number of regions known to play a role within a distributed socioemotional network, scant functional connectivity data exists directly addressing how these regions communicate. Indeed, only one study investigating empathy deficits in a patient with depersonalization disorder (body self-awareness disruption) employed graph-theory analyses, and demonstrated impaired Affective empathy and IA network measurements in the AI, ACC and somatosensory cortex (Sedeño et al. 2014). Although germane, the study only supports an association between these domains during active, task-relevant network configurations. Thus far, no evidence exists regarding this interrelation in a healthy sample during resting state (rsfMRI), despite studies showing that individual differences in complex social processing (i.e. empathy) are reflected in the brain’s intrinsic connectivity networks (ICNs) (Tavor et al. 2016; Smith et al. 2009; Bilevicius et al. 2018; Christov-Moore et al. 2020; Cox et al. 2012).

In addition to connectivity, ICN variability is an often-discounted neuroimaging measurement that may offer complementary information regarding network function and organization. Recent theories consider high variability necessary for the neural system’s adaptability, efficiency and cognitive performance (Dai et al. 2016; Garrett et al. 2010; Garrett, Kovacevic, et al. 2013; McIntosh, Kovacevic, and Itier 2008; Vakorin et al. 2011; Vakorin, Lippé, and McIntosh 2011). Specifically, according to the coordination dynamics theory, networks demonstrating increased variability can flexibly shift through integrative and segregative configurations, maintaining the neural system in balance (Tognoli and Kelso 2014). Thus far, network variability has been predominantly used in clinical populations to investigate alteration in the organization of brain networks (Scarapicchia et al. 2018; Zhang et al. 2020; Good et al. 2020; Kumral et al. 2020) and in healthy populations to investigate brain maturation trajectories (Nomi et al. 2017). To the best of knowledge however, no study has explored the intrinsic network characteristics of connectivity and variability that putatively underlie the socioemotional constructs of empathy and IA in a healthy sample.

Therefore, the present study employs a data-driven approach to explore resting state connectivity and variability related to brain networks underlying Cognitive, Affective empathy and IA. Specifically, we aim to understand whether Cognitive and/or Affective empathy as measured by self-report questionnaires share intrinsic connectivity and/or variability with IA during the viewing of naturalistic stimuli in healthy adults. We hypothesize based on previous literature that 1) Affective empathy will share network connectivity with IA within amygdala, AI and rIFO, given their involvement in the processing of emotion experienced in oneself and vicariously for others (Singer et al. 2004), while 2) Cognitive empathy will share network variability with IA within the rTPJ and precuneus, as these regions are posited to underlie explicit mentalizing (Naughtin et al. 2017; Kovács et al. 2014; Hyde, Aparicio Betancourt, and Simon 2015; Bardi et al. 2017).

## 2 Materials and Methods

### 2.1 Participants and Procedure

26 healthy young adults (m=21.85 y.o./16 female) without a reported history of neurological or psychiatric disorders were recruited for this study **(Table 1)**. All participants were right-handed and had normal or corrected-to-normal vision and hearing. Participants were recruited through on-campus flyers. All participants were paid $20 for their participation. Experimental protocols were approved by University of Louisville’s Institutional Review Board prior to data collection, and written informed consent was obtained from each participant prior to experimental sessions. The study took part on two separate days. On the first day, participants visited the lab to be briefed on the MRI protocol, fill out consent forms and behavioral assessments. On the second day, participants completed the rsfMRI scan at the University of Louisville, School of Medicine.

**Table 1.**
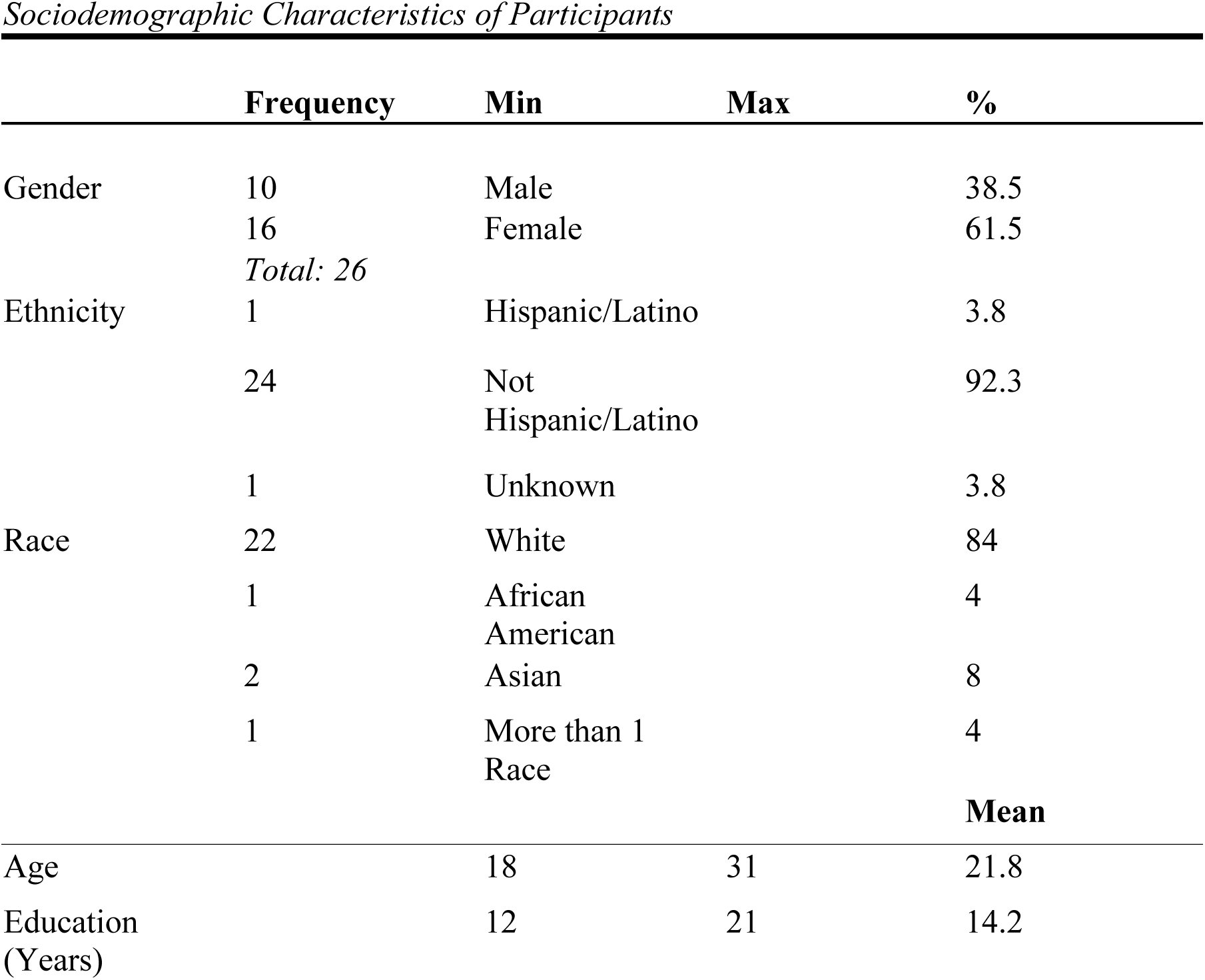

### 2.2 Resting State Scan: Naturalistic Paradigm

The participants watched a 7 min abstract, nonsocial movie titled *Inscapes* previously demonstrated to evoke strong connectivity in networks that resemble rest more than those exhibited during conventional movies (Vanderwal et al. 2015). The movie features a series of technological-looking abstract shapes (**Figure 1**). Participants were told to keep their eyes open and relax while watching and listening to the movie. The stimulus was displayed using E-prime on an Invivo Esys LCD TV monitor at the back of the scanner bore, which was viewed by participants through a mirror on the head-coil. The video is freely available for download from HeadSpace Studios.

**Figure 1:**
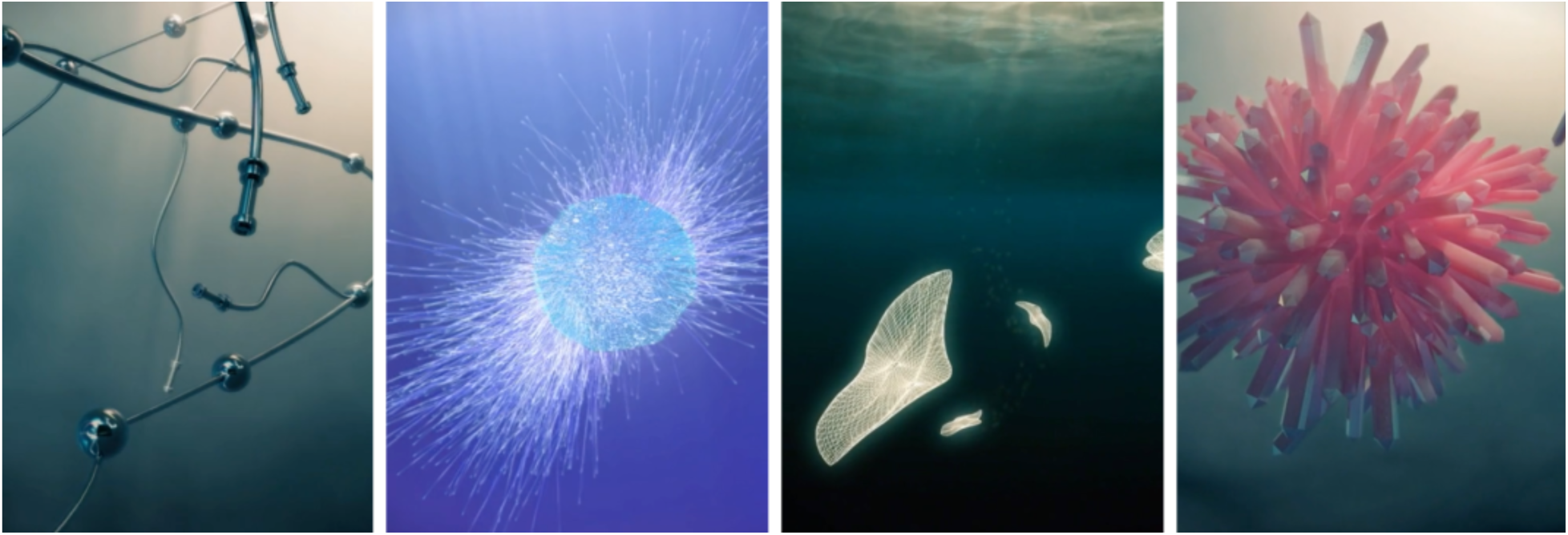
Still-shots from the InScapes Movie.

### 2.3 Behavioral Assessments

#### 2.3.1 Empathy Questionnaire – Interpersonal Reactivity Index (IRI)

Affective and Cognitive empathy was assessed using the Interpersonal Reactivity Index (IRI) (Davis 1983). The IRI consists of 28 items rated on a 5-point scale with the anchors: “does not describe me well” to “describes me very well”. The items are arranged into four subscales with seven items. Each subscale measures a distinct component of empathy: empathic concern (EC) (feelings of compassion and concern for others); personal distress (PD) (feelings of anxiety and discomfort that result from observing another person’s negative experience); perspective taking (PT) (the ability to adopt the perspectives of other people and see things from their point of view); and fantasy subscale (FS) (the tendency to identify with characters in movies, books, or other fictional situations) (Davis 1983). Affective empathy, the ability to infer an agent’s feelings or emotions, was derived from summing the EC and PD subscales. Cognitive empathy, the ability to infer an agent’s beliefs or thoughts, was derived from summing the FS and PT subscales. Total empathy was derived from aggregating Affective and Cognitive empathy scores. All scores were standardized by applying a z-score transformation, and later compared with the MAIA assessment and its subscales (see below) in subsequent analyses.

#### 2.3.2 Multidimensional Assessment of Interoceptive Awareness (MAIA)

The Multidimensional Assessment of Interoceptive Awareness (MAIA) is a 32-item instrument assessing IA: “the conscious perception of sensations from inside the body that create the sense of the physiological condition of the body, such as heart beat, respiration, satiety, and the autonomic nervous system sensations related to emotions” (Mehling et al. 2012). Each statement is rated from 0 (never) to 5 (always) in terms of how often it applies to the participant generally in daily life. The statements are then separated into 8 subscales: Noticing, Non-Distracting, Not-Worrying, Attention-Regulation, Emotional Awareness, Self-Regulation, Body Listening and Trusting, which are in turn aggregated into 5 overall scales used in the present study: Awareness of Body Sensations (Noticing); Emotional Reaction and Attentional Response to Sensations (Not-Distracting, Not-Worrying); Capacity to Regulate Attention (Attention Regulation), Awareness of Mind-Body Integration (Emotional Awareness, Self-Regulation, Body Listening) and Trusting Body Sensations (Trusting). A total Interoceptive Score (MAIA Total) was derived by summing all the aggregate scales. All scores were standardized by applying a z-score transformation, and later compared with the IRI and its subscales (see above) in subsequent analyses. Correlations between behavioral measurements were conducted with the Statistical Package for Social Sciences (Version 25.0.0; SPSS, INC.), and corrected for age, gender and multiple comparisons using false discovery rate *p*<0.05 (FDR) (Benjamini and Hochberg 1995).

### 2.4 MRI Data Acquisition and Preprocessing

All structural MRI images were acquired using a Siemens 3-T Skyra MR scanner. A 20-channel head coil was used for radiofrequency reception. Participants were given earplugs to reduce scanner noise and were additionally given headphones to receive instructions. Foam padding was added to limit motion if additional room remained within the head coil, and a piece of folded tape was placed over the participant’s forehead as a reminder to remain still throughout the scan. Structural images were obtained via a T1-weighted magnetization-prepared rapid gradient-echo sequence (MPRAGE) in 208 sagittal slices. Imaging parameters were as follows: echo time (TE) = 2.26 ms, repetition time (TR) = 1700 ms, flip angle = 9.0°, field of view (FoV) = 204 mm, and voxel size = 0.8 × 0.8 × 0.8 mm. Scan parameters were consistent for all imaging sessions associated with this study. Functional blood oxygenation level-dependent (BOLD) images were collected using multi-band acceleration factor of 3. Imaging parameters were as follows: TE = 29 ms; TR = 2000 ms; flip angle = 62°; FoV = 250 mm; isotropic voxel size = 2.0 mm^3^; 78 interleaved slices, GRAPPA on, Partial Fourier 7/8. Slices were oriented obliquely along the AC–PC line.

All analyses were conducted using the CONN toolbox 19.c (Whitfield-Gabrieli and Nieto-Castanon 2012) based on SPM12 (Penny et al. 2007) in the 2017 version of MATLAB. Spatial preprocessing in the CONN toolbox included realignment, normalization and smoothing (6 mm FWHM Gaussian filter) using SPM12 default parameter settings. Anatomical volumes were segmented into gray matter, white matter and cerebrospinal fluid (CSF) areas, and the resulting masks were eroded to minimize partial volume effects. Physiological and other sources of noise were estimated and regressed out using CompCor (Behzadi et al. 2007), a method that performs principal component analysis to estimate the physiological noise from white matter and cerebrospinal fluid for each participant. Motion from the ART toolbox was also included as a confound regressor. Next, a 0.01–0.10 Hz temporal band-pass filter standard for resting-state connectivity analyses was applied to the time series (Nieto-Castanon 2015). In sum, detrending, outlier censoring, motion regression, and CompCor correction were performed simultaneously in a single first-level regression model, followed by band-pass filtering. These corrections yielded a residual BOLD time course at each voxel that was used for subsequent analyses.

### 2.5 Neuroimaging Analysis

#### 2.5.1 Network Connectivity

Group-level independent component analysis (ICA) using the CONN toolbox was conducted to identify networks of functionally connected brain regions during resting state that may be associated with the IRI and MAIA scores. This involved the application of the fastICA algorithm to volumes concatenated across subject and resting state condition in order to identify independent spatial components (ICs) and back-projection of these components to individual subjects, resulting in maps of regression coefficients representing connectivity between the network and every voxel in the brain (see Calhoun et al., 2001 for details). Forty ICs were identified using spatial overlap of suprathreshold areas (Dice coefficient (Rombouts et al. 1998)), based on CONN’s default network atlas with ROIs characterizing an extended set of classical brain networks: Default Mode Network (4 ROIs), SensoriMotor (2 ROIs), Visual (4 ROIs), Salience / Cingulo-Opercular (7 ROIs), DorsalAttention (4 ROIs), FrontoParietal / Central Executive (4 ROIs), Language (4 ROIs), Cerebellar (2 ROIs) (all ROIs defined from CONN’s ICA analyses of HCP dataset / 497 subjects) (Nieto-Castanon 2014). We chose 40 ICs due to research suggesting ICA results are only affected by the number of ICs when the number is smaller than the number of source signals (Ma et al. 2007), in addition to assuring coverage of a majority of signal variance. Noise components were further identified through visual inspection by authors (TS and BD) (e.g., components largely overlapping CSF), resulting in the exclusion of 4 out of 40 ICs from further consideration. Each of the remaining 36 ICs was subsequently entered in multiple regressions with the IRI and MAIA subscales, aggregate and Total scores. For each network, the resulting statistical maps were cluster thresholded at *p*≤0.05, voxel thresholded at *p*<0.001 (FDR-corrected) using Gaussian Random Field Theory (Worsley et al. 1996), and corrected for age and gender. All coordinates reported below refer to peak activations in anatomical MNI space.

#### 2.5.2 Network Variability

In order to assess network variability and its relationship to either empathy or IA, we regressed each IC’s network variability (calculated in CONN as SD of each IC’s BOLD timeseries: SD_BOLD_ (Nieto-Castanon 2020)) with the IRI and MAIA subscales.

## 3 RESULTS

### 3.1 Behavioral Results

In order to establish the relationship between empathy and IA, all IRI and MAIA subscales, aggregate and Total scores were correlated (**Table 2**).

**Table 2:**
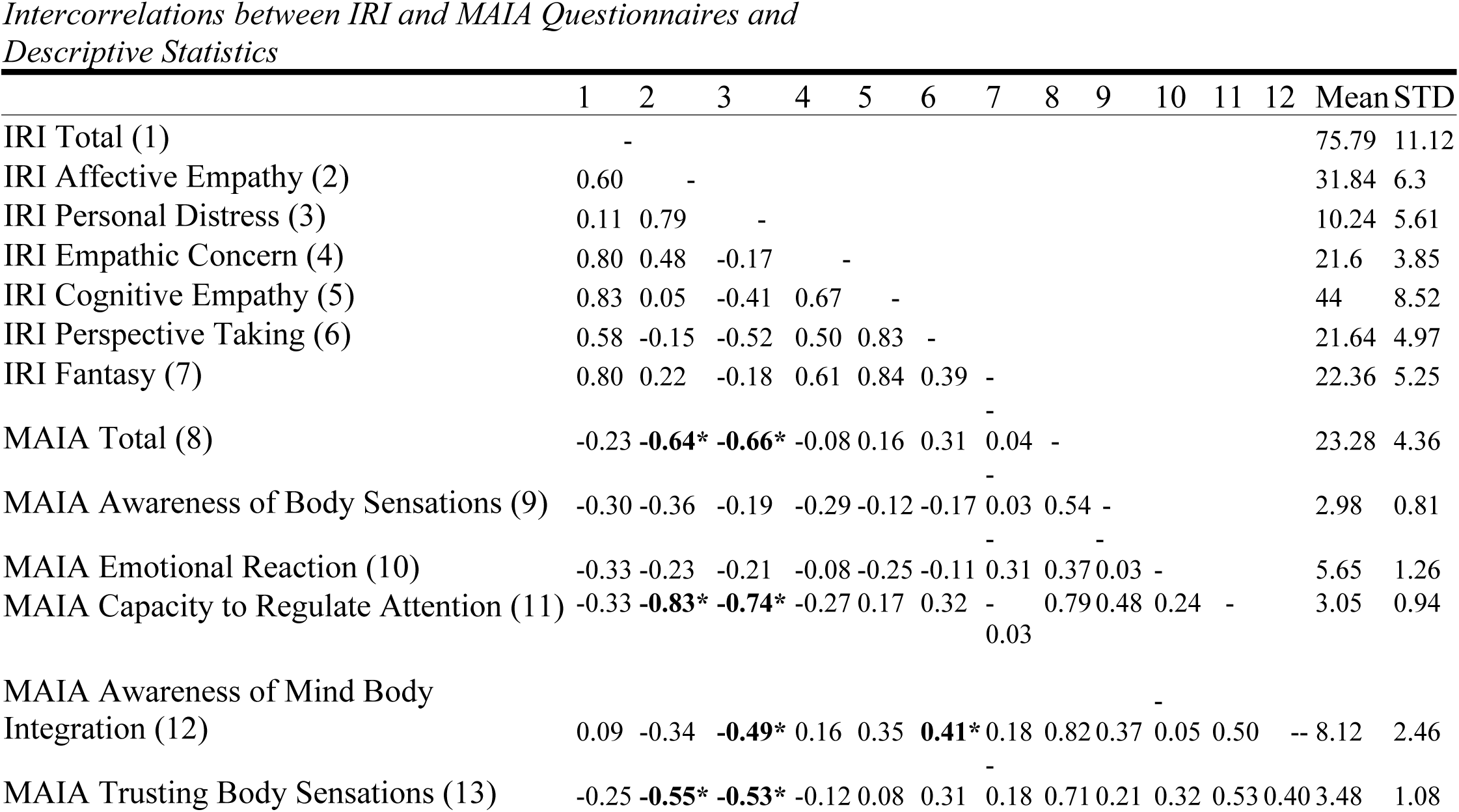
Intercorrelations between IRI and MAIA Subscales and Descriptive Statistics (same subscale correlations not flagged for significance). * represents correlations statistically significant after multiple comparison correction. Uncorrected Mean and STD reported.

#### 3.1.2 Relationship between Affective Empathy and Interoceptive Awareness

Excluding same-subscale correlations, negative relationships were observed between Affective empathy and the following MAIA subscales: Capacity to Regulate Attention (r(26) = - 0.83, *p*<0.01), Trusting Body Sensations (r(26) = −0.55, *p*<0.01) and MAIA Total (r(26) = −0.64, *p*<0.01). Similarly, we observed a negative relationship between the Personal Distress (PD) subscale and Capacity to Regulate Attention (r(26) = −0.74, *p*<0.01), Awareness of Mind Body Integration (r(26) = −0.49, *p*<0.01), Trusting Body Sensations (r(26) = −0.53, *p*<0.01) and MAIA Total (r(26) = −0.66, *p*<0.01). Therefore, we report a negative relationship between Affective empathy and IA (most influenced by the PD scale, since EC exhibited no significant relationship).

#### 3.1.3 Relationship between Cognitive Empathy and Interoceptive Awareness

We observed a positive relationship between Cognitive Empathy and the Awareness of Mind Body Integration subscale, r(26) = 0.35, *p*=0.06, although it did not survive multiple comparison correction. In addition, we observed a significant positive relationship between the IRI Perspective Taking (PT) subscale and the MAIA Awareness of Mind Body Integration subscale, r(26) = 0.41, *p*<0.01. Therefore, we report a positive relationship between Cognitive empathy and IA (most influenced by the PT subscale, since Fantasy exhibited no significant relationship).

Therefore, taken together, our behavioral results show a bidirectional relationship between empathy and IA, depending on the facet of empathy interrogated and mainly driven by the subscales of PD (Affective Empathy) and PT (Cognitive Empathy).

### 3.2 Functional Connectivity Results

#### 3.2.1 Network Connectivity

We observed that within a network comprising right inferior frontal operculum (rIFO), bilateral superior parietal lobule (SPL) and bilateral middle temporal gyrus (MTG), greater connectivity in the rIFO was associated with greater overall empathy as measured by the IRI Total, and with greater IRI Affective empathy. Conversely, lower connectivity in the rIFO was associated with increased overall IA as measured by the MAIA Total and increased MAIA Capacity to Regulate Attention (**Figure 2, Table 3**).

**Table 3:**
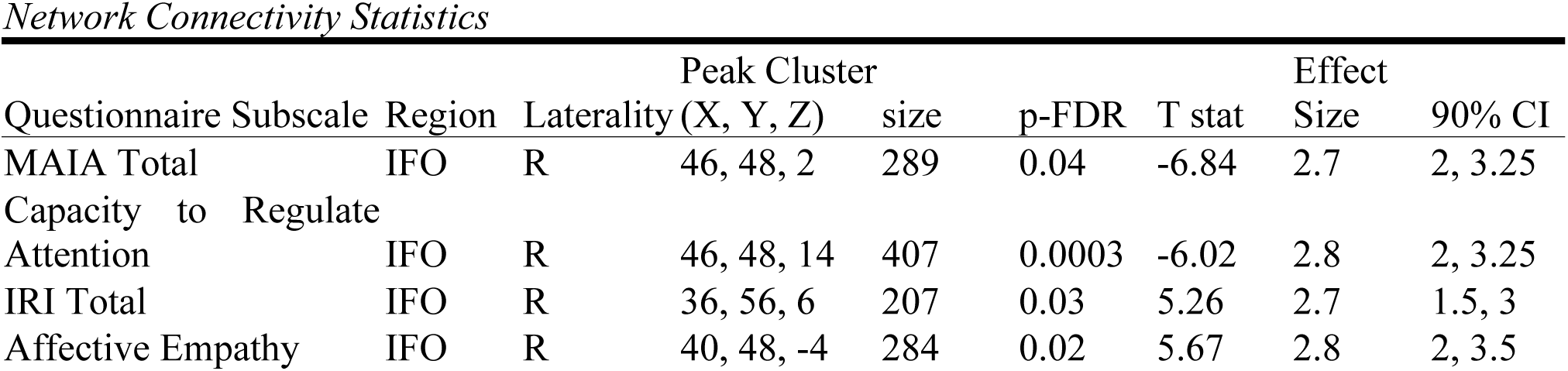
Connectivity Statistics of rIFO cluster within network associated with lower total IA and greater Affective empathy.

| Questionnaire Subscale | Region | Laterality | Peak Cluster |  | p-FDR | T stat | Effect |  |
| --- | --- | --- | --- | --- | --- | --- | --- | --- |
|  |  |  | (X, Y, Z) | size |  |  | Size | 90% CI |
| MAIA Total | IFO | R | 46, 48, 2 | 289 | 0.04 | -6.84 | 2.7 | 2, 3.25 |
| Capacity to Regulate<br>Attention | IFO | R | 46, 48, 14 | 407 | 0.0003 | -6.02 | 2.8 | 2, 3.25 |
| IRI Total | IFO | R | 36, 56, 6 | 207 | 0.03 | 5.26 | 2.7 | 1.5, 3 |
| Affective Empathy | IFO | R | 40, 48, -4 | 284 | 0.02 | 5.67 | 2.8 | 2, 3.5 |

**Figure 2:**
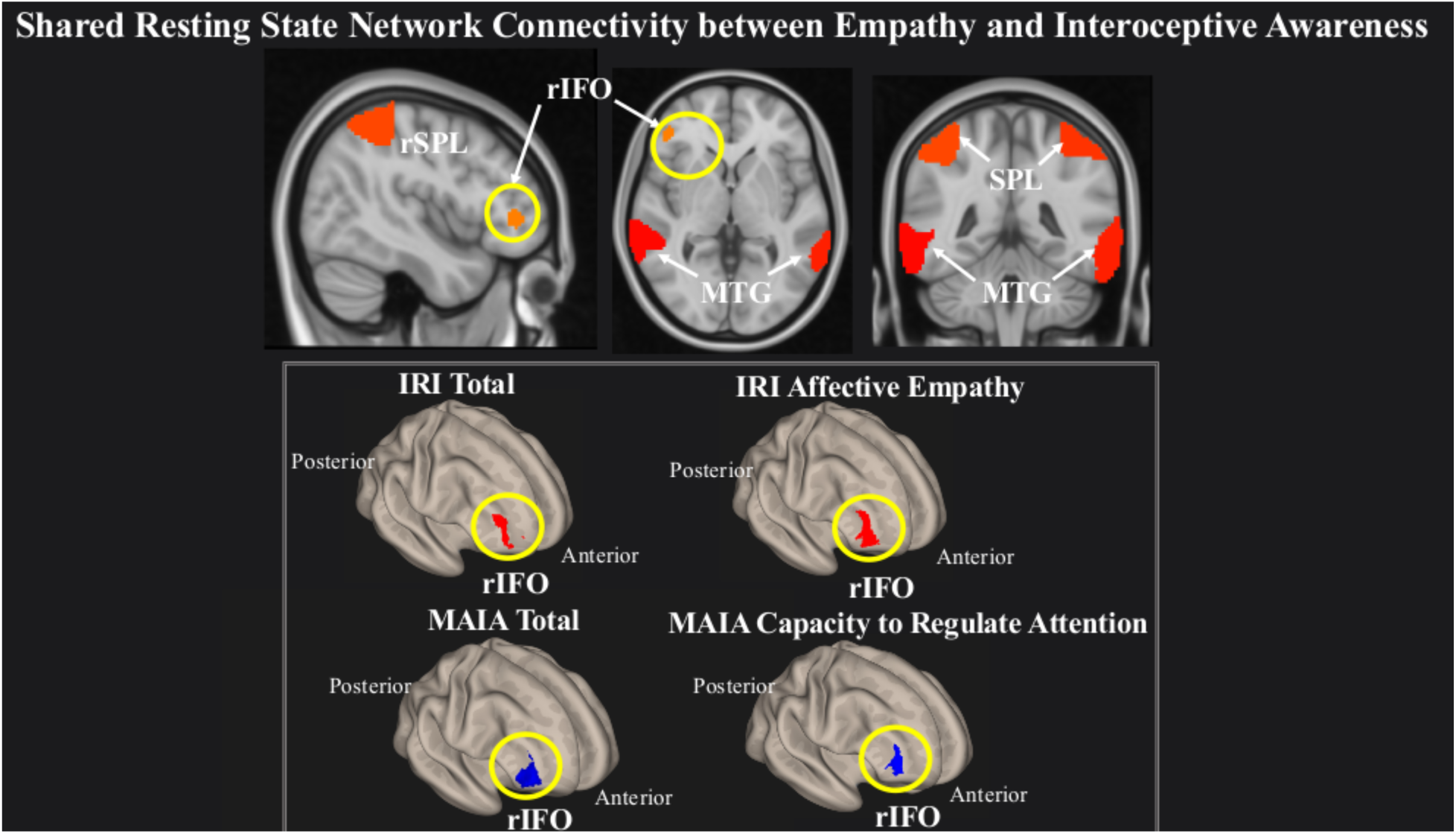
Greater connectivity in the rIFO is associated with lower total IA and greater Affective empathy. Statistical maps are thresholded at FDR-corrected cluster-based *p* < 0.05 after voxel threshold at *p* < 0.001.

#### 3.2.2 Network variability

Network variability analyses revealed that within a network comprising left IFO (L IFO), Cerebellum, and rAI, IRI Personal Distress was negatively related to variability of the network (T(22) = −1.04, *p*=0.02)), while conversely, MAIA Awareness of Body Sensations was positively related (T(22) = 1.48, *p*=0.001)), (**Figure 3**).

**Figure 3:**
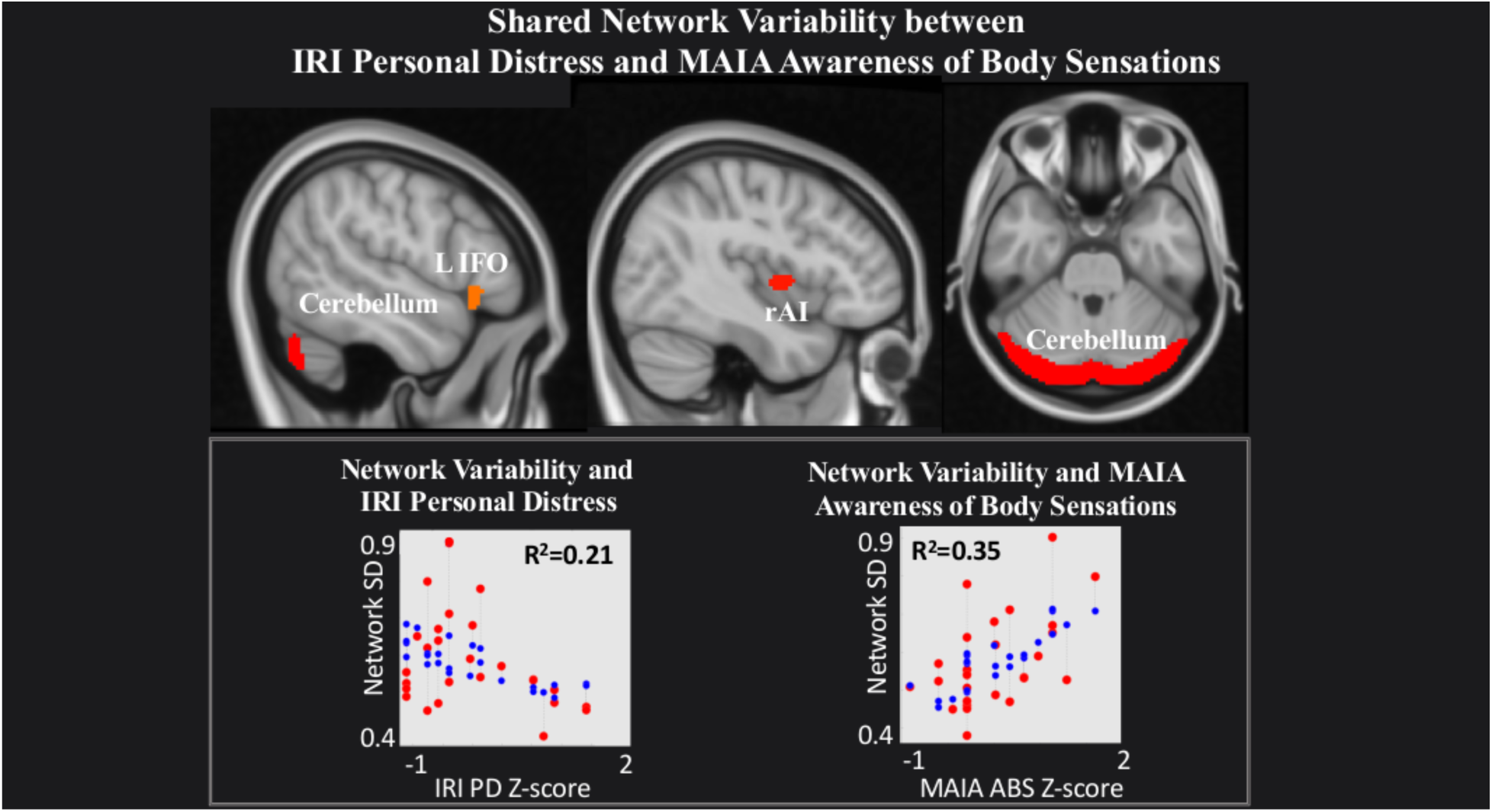
Shared network variability between IRI Personal Distress and MAIA Awareness of Body Sensations. Statistical maps are thresholded at FDR-corrected cluster-based *p* < 0.05 after voxel threshold at *p* < 0.001. *Note:* Red dots in graphs represent observed values, Blue dots fitted values.

Lastly, within a network comprising right Precuneus, rMTG, bilateral supramarginal gyrus (SMG) and rIFO, IRI Perspective Taking (T(22) = 1.45, *p*=0.001) and MAIA Mind Body Integration (T(22) = 2.52, *p*=0.01) were positively related to network variability (**Figure 4**).

**Figure 4:**
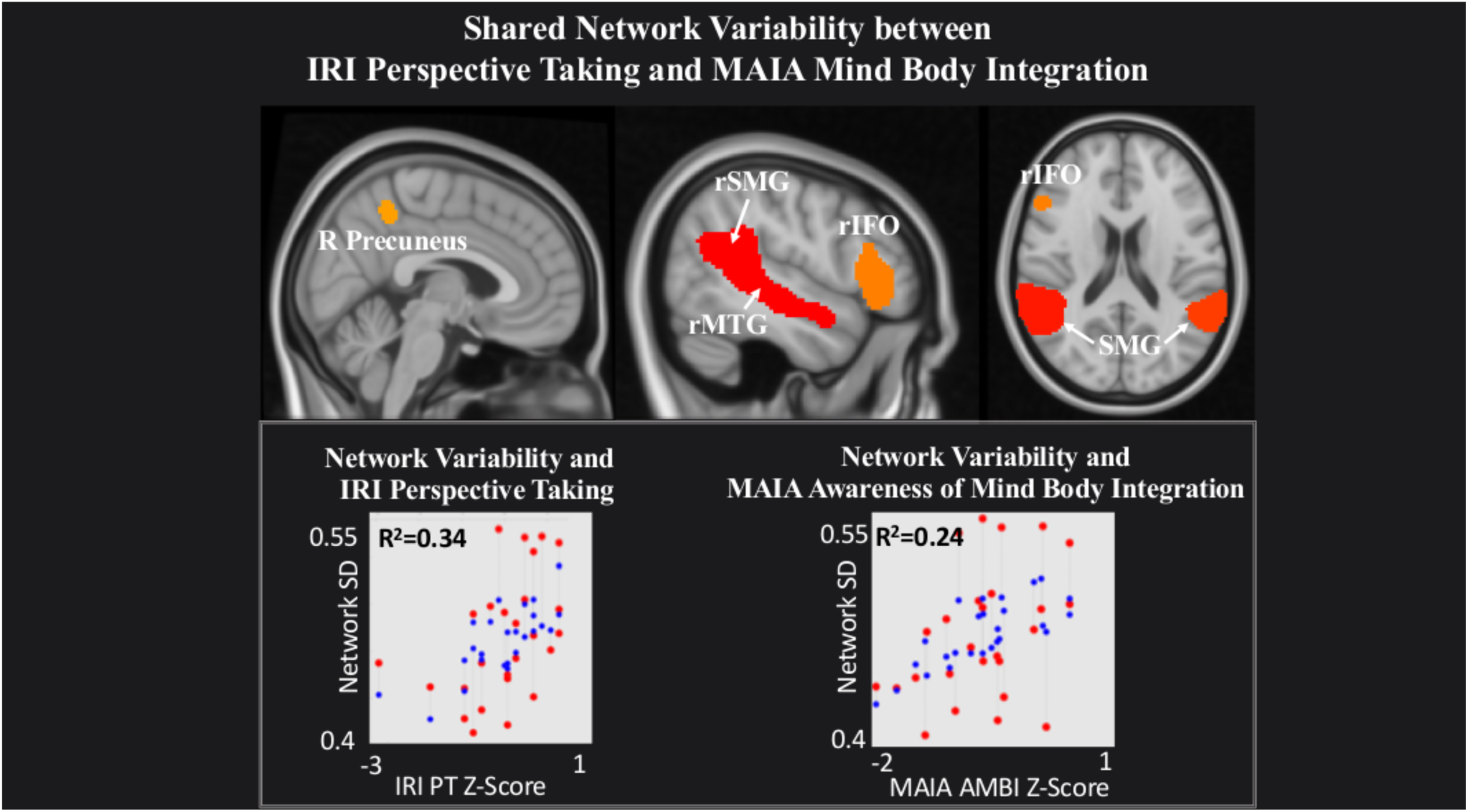
Shared network variability between IRI Perspective Taking and MAIA Mind Body Integration. Statistical maps are thresholded at FDR-corrected cluster-based *p* < 0.05 after voxel threshold at *p* < 0.001. *Note:* Red dots in graphs represent observed values, Blue dots fitted values.

## 4 Discussion

Interoceptive awareness and empathy are core social abilities that enable humans to create meaningful social exchanges by accessing one’s own internal state in order to understand the experiences of others. Our resting state fMRI study employed Independent Component Analysis (ICA) to investigate which empathy facet (Cognitive or Affective) shares network connectivity or variability with interoceptive awareness (IA) while healthy adults viewed naturalistic stimuli. We observed a bidirectional behavioral relationship between empathy and IA, whereby Affective empathy and IA were negatively related, and Cognitive empathy and IA were positively related. This bidirectional link also appeared in the neuroimaging results, such that Affective empathy and IA were inversely related to increased connectivity within the rIFO, and also inversely related to network variability; whereas Cognitive empathy and IA showed only a positive relationship with network variability. Together, these results suggest a double disassociation between empathy and IA depending on the type of empathy interrogated, which is reflected in the brain network’s intrinsic connectivity and variability patterns.

Behaviorally, we observed a negative relationship between the Personal Distress (PD) subscale of the Affective empathy aggregate IRI scale and the total MAIA score, Capacity to Regulate Attention and Trusting Body Sensations subscales. The Capacity to Regulate Attention subscale pertains to various ways of controlling one’s attention towards bodily sensations, as part of an active regulatory process; while Trusting Body Sensations reflects the extent to which one views awareness of bodily sensations as helpful for decision making (Mehling et al. 2012). The Empathic Concern (EC) subscale of Affective empathy exhibited no significant relationship, implying that PD is the dominant subscale of the Affective empathy aggregate scale when relating to the MAIA. This distinction is important, considering that previous data suggests EC motivates individuals to pay attention *to others’* emotions in order to try to comfort them, while conversely, PD drives attention *away from others* in order to reduce the aversive effects for oneself, perhaps as a form of emotion regulation (Zaki et al. 2014). Indeed, Decety & Jackson, 2004 proposed that PD may arise from the failure of applying sufficient self-regulatory control over the shared emotional state. In line with previous studies, we report an inverse relationship between PD and a an attention regulation measure – MAIA’s Capacity to Regulate Attention subscale. Together with the Trusting Body Sensations subscale, our findings suggest the increased ability to regulate internal attention and rely on this discrete information may be linked to a decrease in the discomfort experienced while witnessing another’s distress.

Furthermore, we found a positive relationship between the Perspective Taking (PT) subscale of the Cognitive empathy aggregate IRI scale and the Awareness of Mind Body Integration of the MAIA. This MAIA subscale represents the integration of several higher level cognitive processes necessary for socially relevant goal-directed behavior (i.e. executive functions (Pribram 1973)) including: emotional awareness, self-regulation of emotions, and the ability to feel a sense of an embodied self, that is – “a sense of the interconnectedness of mental, emotional, and physical processes as opposed to a disembodied sense of alienation from one’s body” (Mehling et al. 2012). Thus, our results support previous findings suggesting Cognitive empathy, and in particular PT, is related to a wide array of executive function skills such as working memory, inhibitory control and cognitive flexibility (Yan et al. 2020; Aliakbari et al. 2013; Healey and Grossman 2018). Taken together with the aforementioned negative relationship between PD and IA, these behavioral results suggest a bidirectional association between empathy and IA, contingent on the type of empathy interrogated. To wit, directing attention towards internal bodily sensations may relieve vicarious emotional pain but flexibly employing cognitive-control skills may increase the ability to take the perspective of another in pain.

Our connectivity results provide further support for this inverse relationship. Within a network of brain regions previously shown to underlie attentional processing (superior parietal lobule (SPL), medial temporal gyrus (MTG) and right inferior frontal operculum (rIFO) (Corbetta and Shulman 2002; Wu et al. 2016; Perrett et al. 1992; Perrett, Rolls, and Caan 1982; Perrett et al. 1985), we observed that increased connectivity in the rIFO was associated with increased overall empathy (total IRI score) and the Affective empathy aggregate scale on one hand, but reduced overall IA (total MAIA score) and Capacity to Regulate Attention on the other hand. Previous studies investigating both personal (Craig 2009; Johnson 2001; Damasio 2005; Critchley 2005; Gray et al. 2007) and vicarious emotional experience (Singer et al. 2004; Jabbi, Swart, and Keysers 2007) show the consistent activation of the anterior insula (AI) and frontal operculum, and therefore the IFO is thought of as a continuum between these two structures (Jabbi, Swart, and Keysers 2007; Wicker et al. 2003). Because Affective empathy was driven by the PD subscale, increased connectivity within the rIFO in the present study may relate to intensified personal suffering from witnessing another’s distress, but decreased awareness of one’s own body sensations, perhaps due to allocating attention externally (for example, away from oneself and toward another’s distress). Our findings reflect previous results showing neural overlap of interoception and empathy in this region (Ernst et al. 2013; Adolfi et al. 2017), and extend these findings by providing intrinsic connectivity evidence of a double dissociation between empathy and IA.

Our network variability results offer a complementary perspective that further supports this bidirectional relationship. We observed increased scores on the MAIA Awareness of Body Sensations subscale and decreased scores on the IRI PD subscale was associated with increased variability of brain regions previously shown to underlie processing and integration of visceral information (i.e., Cerebellum, L IFO, L AI) (J. D. Schmahmann 2001; Baumann and Mattingley 2012; Schienle and Scharmüller 2013; Adamaszek et al. 2017; Bogg and Lasecki 2015; Terasawa, Fukushima, and Umeda 2013). Despite the prevailing focus on the AI as a hub for processing body sensations (Terasawa, Fukushima, and Umeda 2013; Kuehn et al. 2016; Pollatos, Gramann, and Schandry 2007; Singer, Critchley, and Preuschoff 2009), additional brain regions are also commonly implicated in interoceptive experience. For example, fMRI studies identify the involvement of the IFO and cerebellum, reinforcing the notion that interoceptive processing (and perhaps especially nociceptive information) may occur through multiple neural pathways (Rapps et al. 2008; Peiffer et al. 2008; Garcia-Larrea 2012). In the same vein, observing distress in others without actually experiencing it may rely on high-order cognitive functions to access minor changes in physical state, as a tool to modulate negative stimulus input (Preckel, Kanske, and Singer 2018). The implication of the cerebellum in a shared network underlying both PD and interoceptive processing is not surprising, since the cerebellum serves as an integral node in the distributed cortical–subcortical neural circuitry supporting an array of social cognitive operations (Jeremy D. Schmahmann and Pandya 1995). Taken together, we suggest variability of this network of brain regions underlying interoceptive experience may reflect increased flexibility of the network to process internal sensations, but less adaptability to integrate vicarious emotions arising from witnessing another’s distress.

In addition, we observed a positive relationship between variability of both the IRI PT subscale and the MAIA Awareness of Mind Body Integration subscale within a network of brain regions previously associated with the process of mentalizing – the precuneus, rIFO, SMG and MTG (Mar 2011; Northoff et al. 2006; Schurz et al. 2014; Spreng, Mar, and Kim 2009; Vogeley et al. 2001). Mentalizing, also referred to as Theory of Mind (TOM), signifies the ability to attribute mental states to another individual, allowing the observer to predict intent and direct their behavior appropriately (Frith, Morton, and Leslie 1991; Premack and Woodruff 1978). Researchers agree that mentalizing differs from the vicarious sharing of emotion in its psychological complexity, combining observation, memory, knowledge, and reasoning to provide insight into the thoughts and feelings of others (van der Heiden et al. 2013; Decety and Jackson 2004). Therefore, its connection with MAIA’s Awareness of Mind Body Integration scale is not surprising, since both concepts require not only affective experience, but also comprehension and integration of that particular state of mind within one’s own emotional schema. We therefore suggest that increased variability of brain regions underlying a mentalizing network may point to enhanced network flexibility subserving not only a better ability to take another’s perspective, but also an improved sense of interconnectedness between one’s own mind and body.

Lastly, our data shows an interesting convergence of empathy and IA within the IFO. Research suggests the IFO serves as both a sensory-cognitive integration area and a control node of the ventral attention network (Craig 2009; Corbetta and Shulman 2002), conjectured to maintain goal-related information online until a decision is reached (Tops and De Jong 2006; Tops et al. 2010). Moreover, recent evidence suggests a hemispheric specialization of the IFO related to reactive/proactive goal maintenance (Tops et al. 2010). On one hand, the rIFO may facilitate immediate somatosensory processing and attentional shifting whilst a response is ongoing (reactive) (Tops et al. 2010; Hampshire et al. 2009; Higo et al. 2011; Nelson et al. 2010), through its connections to rostral ACC, superior temporal gyrus (STG) and occipital cortex (Cauda et al. 2011). On the other hand, the lIFO may exert top-down control whilst preparing a response (proactive) (Tops et al. 2010), through its connections to dorsolateral prefrontal cortex and bilateral supplementary motor area (Cauda et al. 2011). Taking this evidence into consideration, we speculate the positive association between connectivity within the rIFO and Affective empathy indicates a propensity in the highly empathic individual to shift attention toward salient cues in their environment (for example, another individual in distress). In contrast, the negative relationship between connectivity within the rIFO and IA may indicate an inability to redirect attention toward external salient cues, and therefore may lead to increased awareness of internal sensations. Our network variability findings offer complementary evidence regarding the role of the IFO in socioemotional processes. We show that increased network flexibility within an interoceptive experience network (comprised of lIFO, L AI, cerebellum) is linked to increased Awareness of Body Sensations as well as decreased PD. We suggest the lIFO plays a crucial part in this network’s ability to modulate attention from one’s own internal sensations to discomfort arising from witnessing another’s distress, perhaps in an effort to plan an appropriate emotional response. In the same vein, we show enhanced network flexibility within a mentalizing network (comprised of rIFO, precuneus, SMG, MTG) is related to both better PT and increased Mind Body Integration. These relationships may illustrate that heightened ability to determine intent and integrate sensory information into one’s own emotional schema could relate to flexibly shifting attention towards the target of interest (either self or other). In sum, our data suggests the IFO serves as an internal/external attention modulator and thus may play a critical role in switching attention from one’s own body sensations (IA) to another’s (Affective and/or Cognitive empathy). Our study’s findings should be considered along with its limitations. The definition of network variability has been inconsistent across previous studies, (e.g., amplitude, variance, standard deviation, mean squared successive difference; for a review, see (Garrett, Samanez-Larkin, et al. 2013) with considerable range in the methodology used to derive them. Therefore, implementation of network variability as a consistently used neuroimaging measure will require increased efforts toward methodological standardization. It is also important to note that due to the nature of the analyses used, the findings of this study do not represent causal relationships. That is, the results represent a correlational relationship between a questionnaire-based measure of IA or empathy and functional connectivity and/or variability. Our sample was unfortunately not large enough for a gender-specific analysis, as evidence suggests there are differences in the capacity for empathy between males and females (Christov-Moore et al. 2014). Future research should be conducted in this regard. Similarly, in using an undergraduate sample, the generalizability of these findings is limited.

In conclusion, the current research provides novel information about the relationship between internal body sensation awareness and empathy. In contrast to previous studies which used task-based fMRI to assess the neurobiology of these two constructs separately, we used a data-driven resting state approach to test whether distinct empathy facets share network characteristics (connectivity and/or variability) with IA. We demonstrate a bidirectional relationship between empathy and IA, depending on the type of empathy investigated. Specifically, Affective empathy and IA share network connectivity and variability, while Cognitive empathy and IA only share variability. In regard to Affective empathy, increased vicarious emotional experience and decreased IA were associated with increased connectivity within the rIFO of a larger attention network; while increased IA and reduced personal discomfort arising from witnessing another’s distress were related to increased network flexibility within an internal sensation network. Concerning Cognitive empathy, perspective-taking ability and a sense of mind-body connectedness related to increased network flexibility between brain regions subserving a mentalizing network. We also suggest the role of the IFO as an internal/external attention modulator that may play a critical role in switching attention from one’s own body sensations (IA) to another’s (Affective and/or Cognitive empathy). Overall, we show that the ability to feel and understand another’s emotional state is related to one’s own awareness of internal body changes, and that this relationship is not task-dependent, but is reflected in the brain’s resting state neuroarchitecture. Methodologically, this work highlights the importance of utilizing network variability alongside functional connectivity as an important complementary route into understanding neurological phenomena. Our results hold promise in aiding diagnosis of clinical disorders characterized by IA and empathy deficits such as the autism spectrum disorders (ASD), where participants may be unable to complete tasks or questionnaires due to the severity of their socioemotional symptoms.

## Author Contributions

**Teodora Stoica**: Data Curation, Investigation, Project administration, Formal analysis, Methodology, Visualization, Writing – original draft **Brendan Depue**: Resources, Supervision, Writing – Review and editing

## Acknowledgements

Many thanks to Dr. Nick Hindy for invaluable advice regarding writing.

## Data Availability Statement

Raw data and behavioral variables of interest for the empirical results presented here can be found on the Mendeley Data Repository at http://dx.doi.org/10.17632/fmch6sj4sw.1

